## Supplementary Materials for "Self-cleaving 2A peptides allow for expression of multiple genes in *Dictyostelium discoideum*"

### Supplementary Information

| Construct | DNA sequence |
|---|---|
| linker | GGTTCAGCAGGTTCAGCAGCAGGTTCAGGTGAATTT |
| P2A | GGATCAGGAGCAACCAATTTCAGTTTGCTTAAACAAGCAGGTGATGTTGAGGAGAATCCAGGTCCT |
| T2A | GGAAGTGGAGAGGGAAGAGGTAGTTTGTTGACTTGTGGTGATGTAGAGGAGAATCCTGGACCA |
| F2A | GGTTCAGGAGTTAAACAAACCTTAAATTTTGATTTATTAAAGTTAGCTGGAGATGTAGAATCTAATCCTGGACCT |
| E2A | GGTAGTGGACAATGTACAAATTACGCATTGTTGAAGTTAGCTGGTGATGTAGAATCAAATCCAGGTCCT |

**Table S1. DNA sequences of 2A viral peptides**

| Construct | Backbone | Strain | Selection | Main Fig(s) | Supp. Fig |
|---|---|---|---|---|---|
| mNeonGreen_mScarlet-I dual-transcription unit | pDGB Ω1Neo (11) | AX4 | G418 (15 µg/mL) | 2, 3 | S2 |
| mScarlet-I_mNeonGreen dual-transcription unit | pDGB Ω1Neo (11) | AX4 | G418 (15 µg/mL) | 2, 3 | S3 |
| mNeonGreen-linker-mScarlet-I fusion protein | pDGB Ω1Neo (11) | AX4 | G418 (15 µg/mL) | 2, 3 | S4 |
| mScarlet-I-linker-mNeonGreen fusion protein | pDGB Ω1Neo (11) | AX4 | G418 (15 µg/mL) | 2, 3 | S5 |
| mNeonGreen-P2A-mScarlet-I | pDGB Ω1Neo (11) | AX4 | G418 (15 µg/mL) | 2, 3 | S6 |
| mScarlet-I-P2A-mNeonGreen | pDGB Ω1Neo (11) | AX4 | G418 (15 µg/mL) | 2, 3 | S7 |
| mNeonGreen-T2A-mScarlet-I | pDGB Ω1Neo (11) | AX4 | G418 (15 µg/mL) | 2, 3 | S8 |
| mScarlet-I-T2A-mNeonGreen | pDGB Ω1Neo (11) | AX4 | G418 (15 µg/mL) | 2, 3 | S9 |
| mNeonGreen_HygroR dual-transcription unit | pDMDC (this study) | AX4 | Hygro (100 µg/mL) | 4 | S11 |
| mNeonGreen_HygroR dual-transcription unit | pDMDC (this study) | AX4 | Hygro (150 µg/mL) | 4 | S12 |
| mNeonGreen_HygroR dual-transcription unit | pDMDC (this study) | AX4 | Hygro (200 µg/mL) | 4 | S13 |
| mNeonGreen_HygroR dual-transcription unit | pDMDC (this study) | NC28.1 | Hygro (400 µg/mL) | 4 | S17 |
| mNeonGreen_HygroR dual-transcription unit | pDMDC (this study) | NC28.1 | Hygro (600 µg/mL) | 4 | S18 |
| mNeonGreen_HygroR dual-transcription unit | pDMDC (this study) | NC28.1 | Hygro (800 µg/mL) | 4 | S19 |
| mNeonGreen-P2A-HygroR | pDMP2A (this study) | AX4 | Hygro (100 µg/mL) | 4 | S14 |
| mNeonGreen-P2A-HygroR | pDMP2A (this study) | AX4 | Hygro (150 µg/mL) | 4 | S15 |
| mNeonGreen-P2A-HygroR | pDMP2A (this study) | AX4 | Hygro (200 µg/mL) | 4 | S16 |
| mNeonGreen-P2A-HygroR | pDMP2A (this study) | NC28.1 | Hygro (400 µg/mL) | 4 | S20 |
| mNeonGreen-P2A-HygroR | pDMP2A (this study) | NC28.1 | Hygro (600 µg/mL) | 4 | S21 |
| mNeonGreen-P2A-HygroR | pDMP2A (this study) | NC28.1 | Hygro (800 µg/mL) | 4 | S22 |

**Table S2. Table of Flow Experiments**

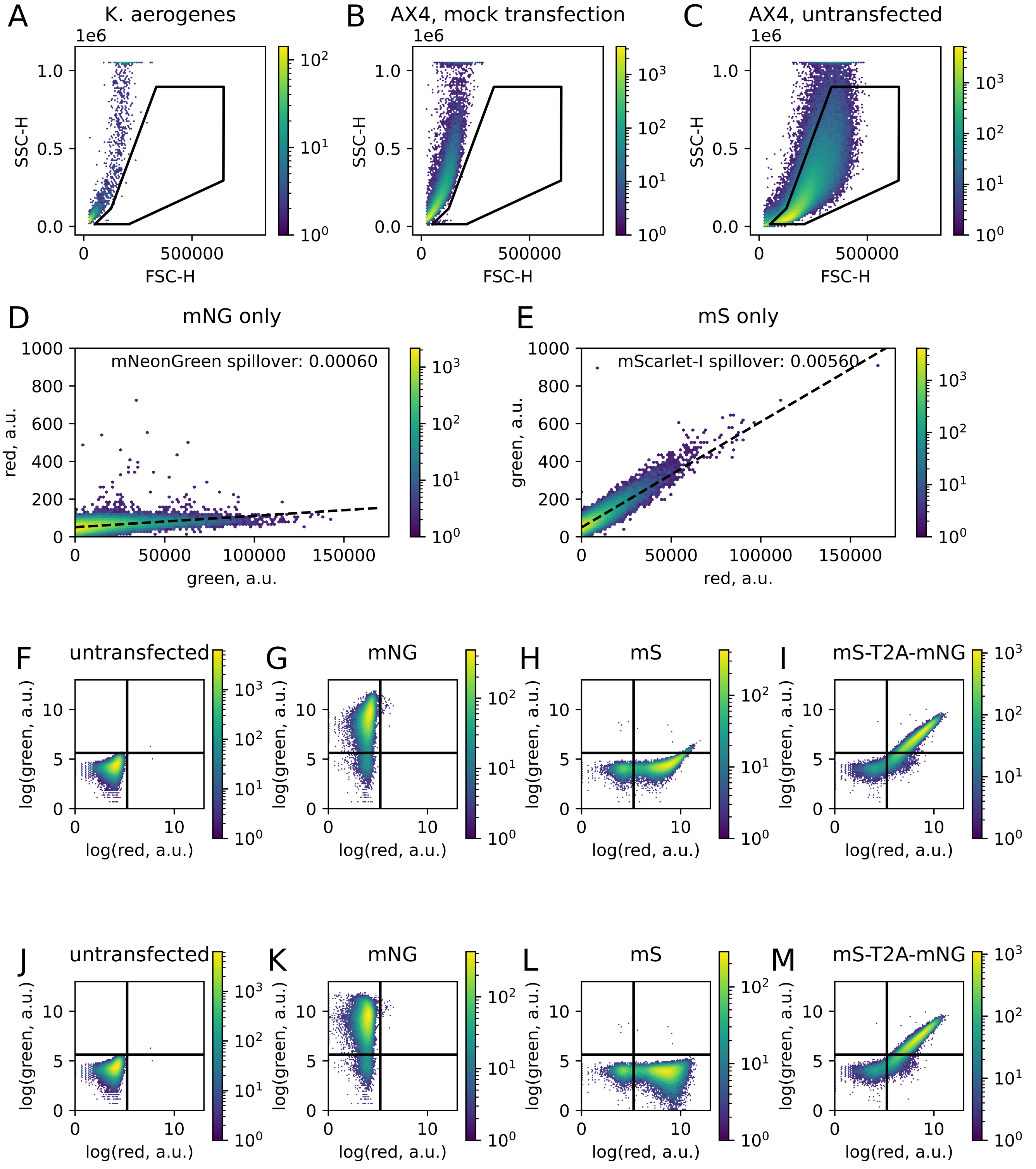


**Fig. S1. Gating and spillover compensation for flow cytometry. A-C.** Forward scatter and side scatter profile of *K. aerogenes* used as a food source for cells post-transfection (A), AX4 cells mock transfected without DNA and incubated with 15 µg/ml G418 for three days (B), and AX4 harvested from a lawn of *Klebsiella aerogenes* on a SM agar plate (C). The black polygon represents the gate separating live Dicty from bacteria, dead cells, or other debris, and only events falling inside the polygon gate were considered for downstream analysis. **D-E.** Fluorescence values for AX4 cells transfected with mNeonGreen only (D) or mScarlet-I only (E). Spillover coefficients for each fluorescent protein were determined by linear fitting. **F-I.** Log-transformed uncompensated fluorescence values for AX4 cells that are untransfected (F), transfected with mNeonGreen only (G), mScarlet-I only (H), or with a co-expression construct (I). The horizontal and vertical lines represent the fluorescence thresholds for green and red, respectively. **J-M.** Fluorescence values from (F-I) after application of the appropriate spillover compensations. Note that J is identical to F because no compensation was required.

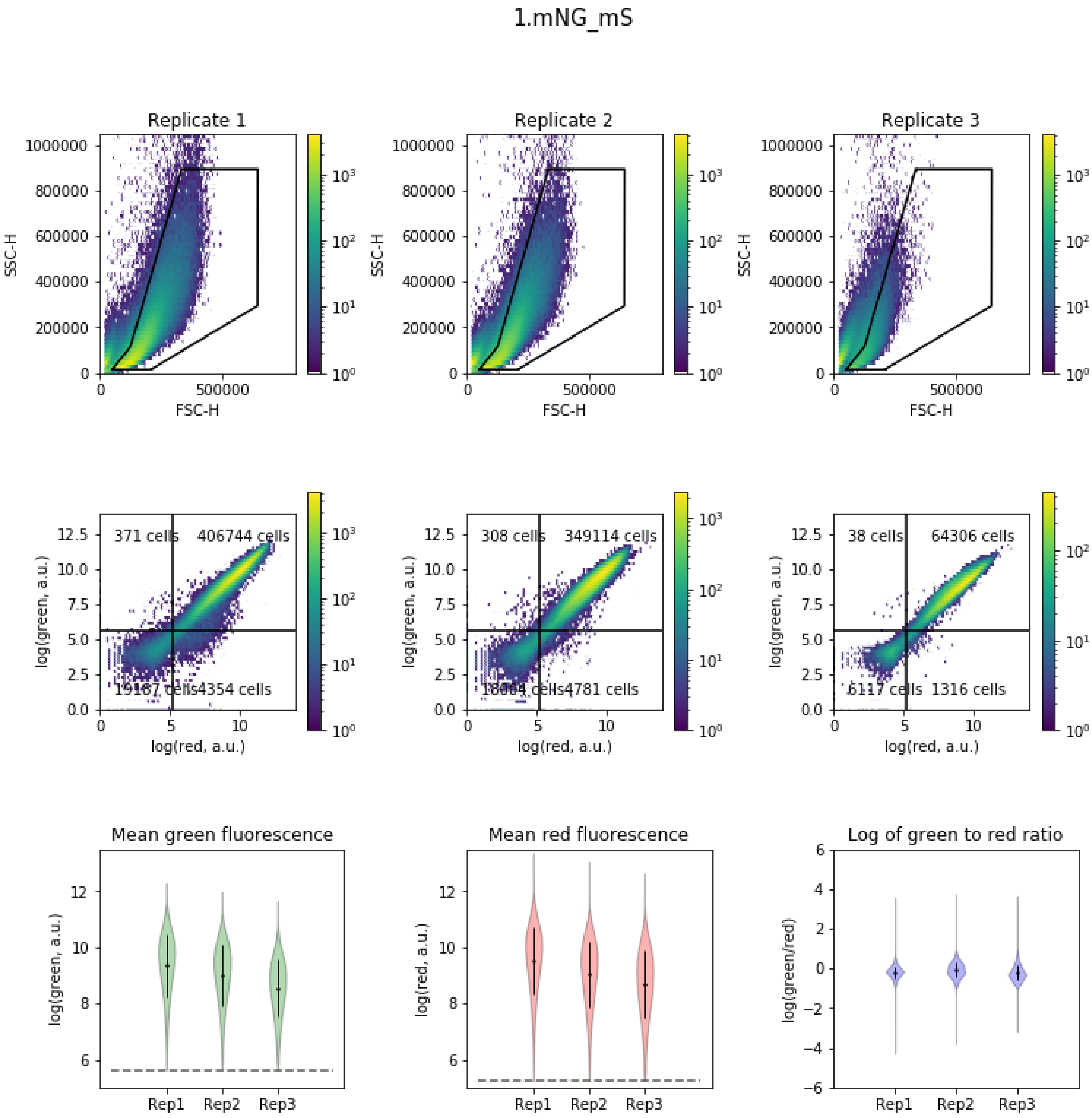


**Fig. S2. Flow cytometry data of AX4 cells transfected with dual transcription cassettes for mNeonGreen and mScarlet-I.Top.** Forward scatter and side scatter profiles of all three replicates. Density map color indicates event count. Events inside the polygon, defined as shown in S1, were considered live cells. **Middle.** Green and red fluorescence values for live cells for each replicate. Displayed numbers indicate total event count in a given quadrant. Cells in the upper right quadrant were retained for further analysis. **Bottom.** Mean fluorescence values and ratio of green to red fluorescence for each replicate. For each violin plot of single-cell values, the overlaid dot represents mean of the distribution and the error bars represent the standard deviation. Dashed lines represent fluorescence thresholds.

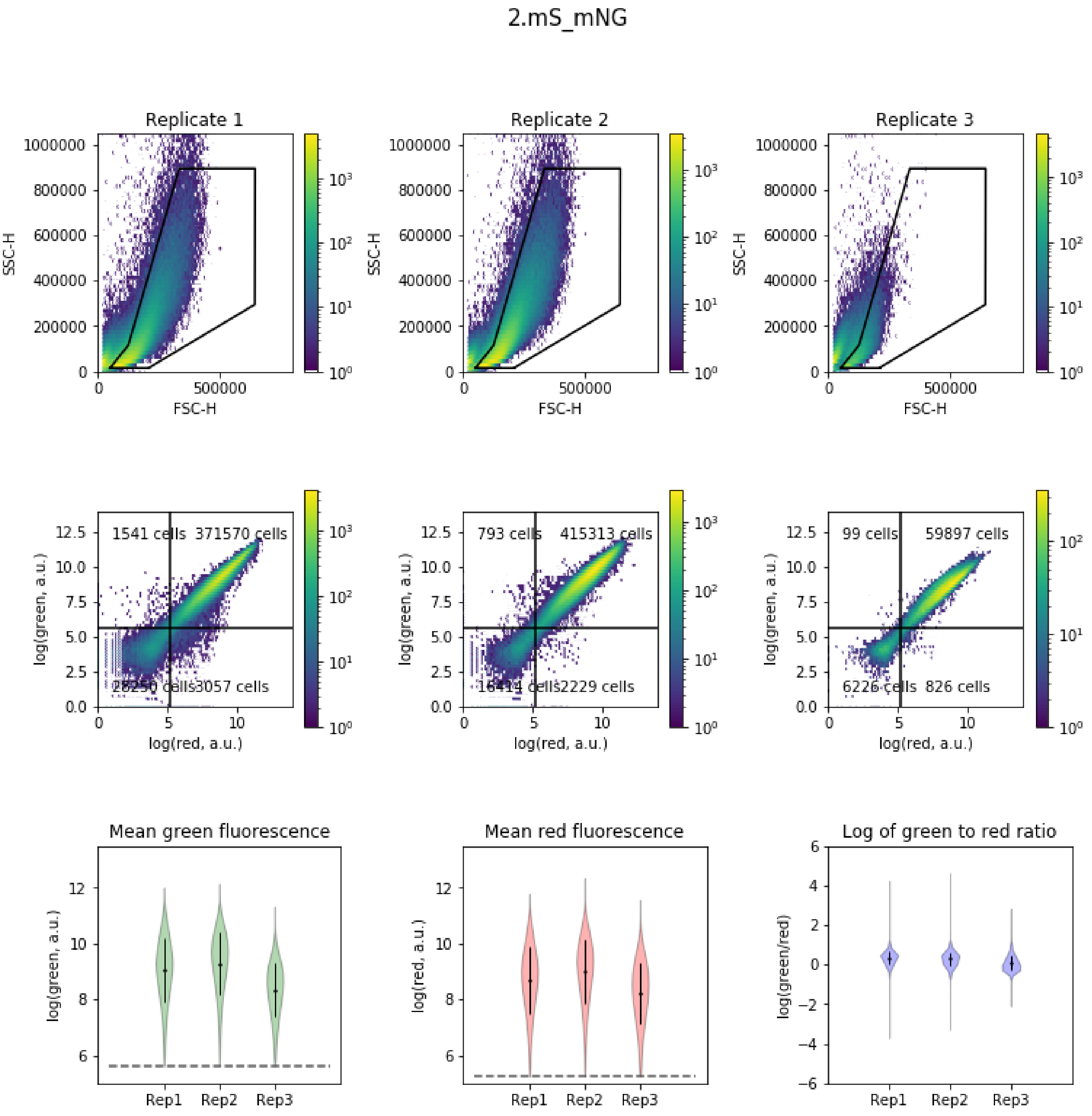


**Fig. S3. Flow cytometry data of AX4 cells transfected with dual transcription cassettes for mScarlet-I and mNeonGreen.Top.** Forward scatter and side scatter profiles of all three replicates. Density map color indicates event count. Events inside the polygon, defined as shown in S1, were considered live cells. **Middle.** Green and red fluorescence values for live cells for each replicate. Displayed numbers indicate total event count in a given quadrant. Cells in the upper right quadrant were retained for further analysis. **Bottom.** Mean fluorescence values and ratio of green to red fluorescence for each replicate. For each violin plot of single-cell values, the overlaid dot represents mean of the distribution and the error bars represent the standard deviation. Dashed lines represent fluorescence thresholds.

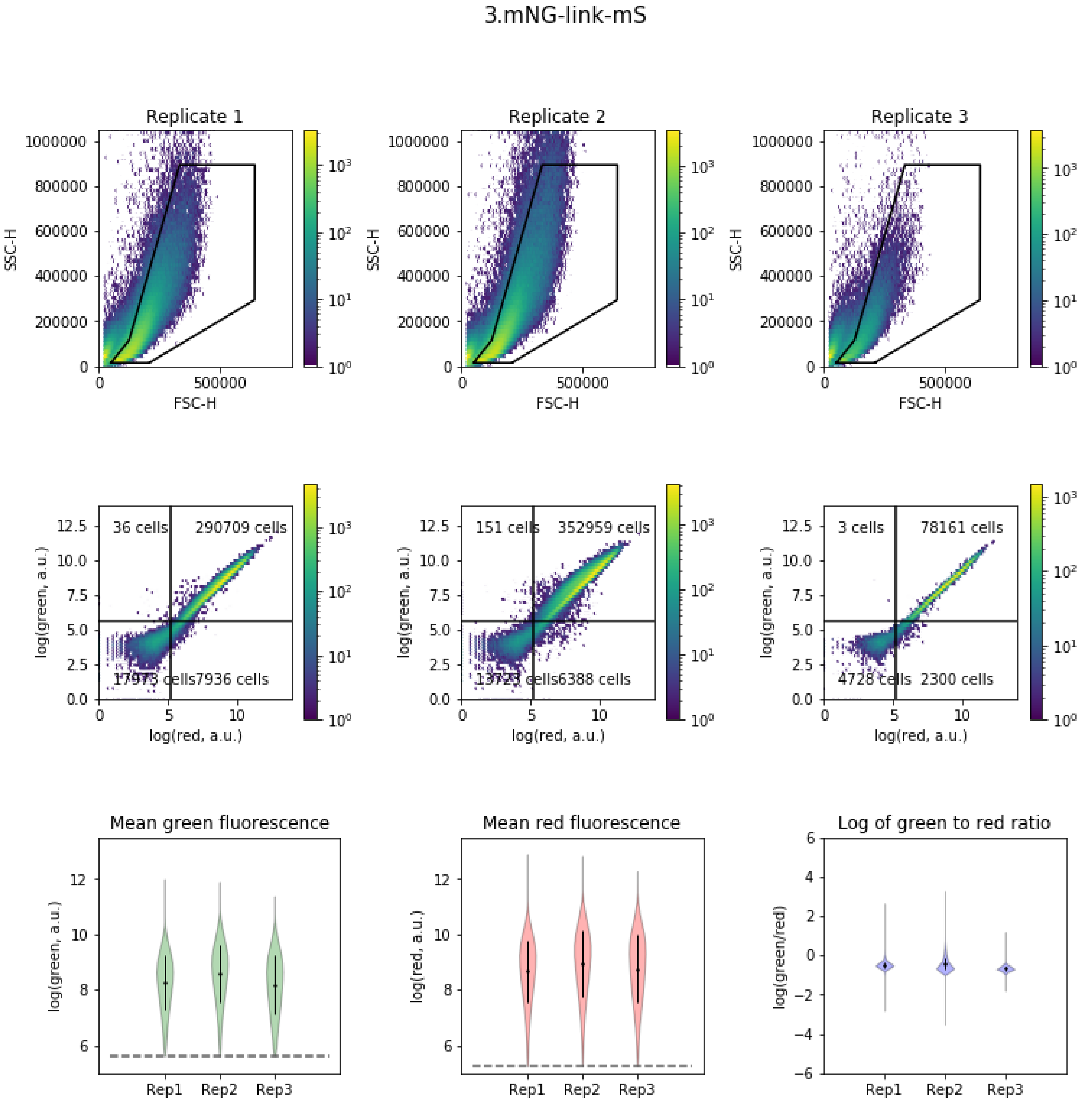


**Fig. S4. Flow cytometry data of AX4 cells transfected with mNeonGreen-linker-mScarlet-I.Top.** Forward scatter and side scatter profiles of all three replicates. Density map color indicates event count. Events inside the polygon, defined as shown in S1, were considered live cells. **Middle.** Green and red fluorescence values for live cells for each replicate. Displayed numbers indicate total event count in a given quadrant. Cells in the upper right quadrant were retained for further analysis. **Bottom.** Mean fluorescence values and ratio of green to red fluorescence for each replicate. For each violin plot of single-cell values, the overlaid dot represents mean of the distribution and the error bars represent the standard deviation. Dashed lines represent fluorescence thresholds.

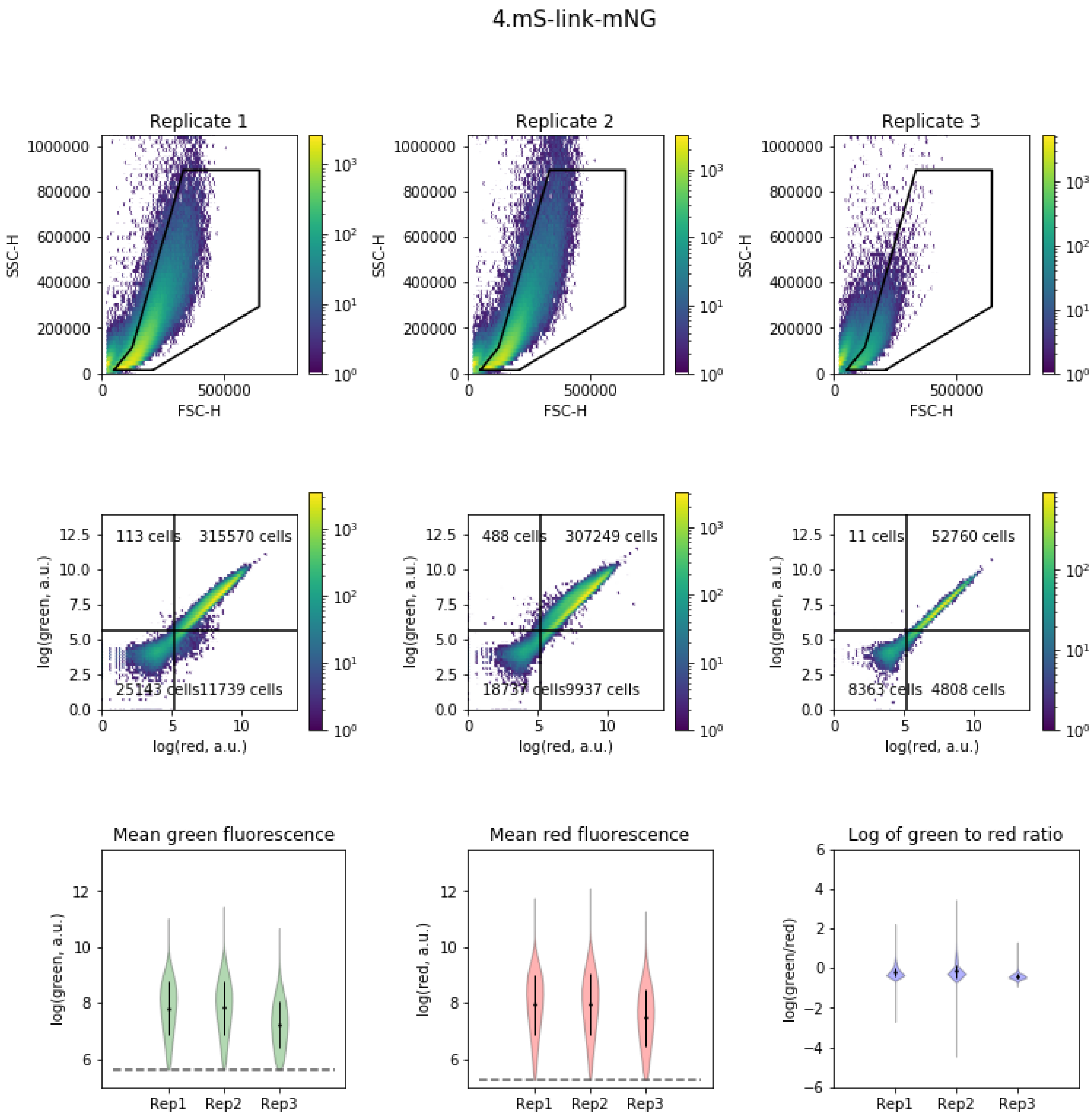


**Fig. S5. Flow cytometry data of AX4 cells transfected with mScarlet-I-linker-mNeonGreen.Top.** Forward scatter and side scatter profiles of all three replicates. Density map color indicates event count. Events inside the polygon, defined as shown in S1, were considered live cells. **Middle.** Green and red fluorescence values for live cells for each replicate. Displayed numbers indicate total event count in a given quadrant. Cells in the upper right quadrant were retained for further analysis. **Bottom.** Mean fluorescence values and ratio of green to red fluorescence for each replicate. For each violin plot of single-cell values, the overlaid dot represents mean of the distribution and the error bars represent the standard deviation. Dashed lines represent fluorescence thresholds.

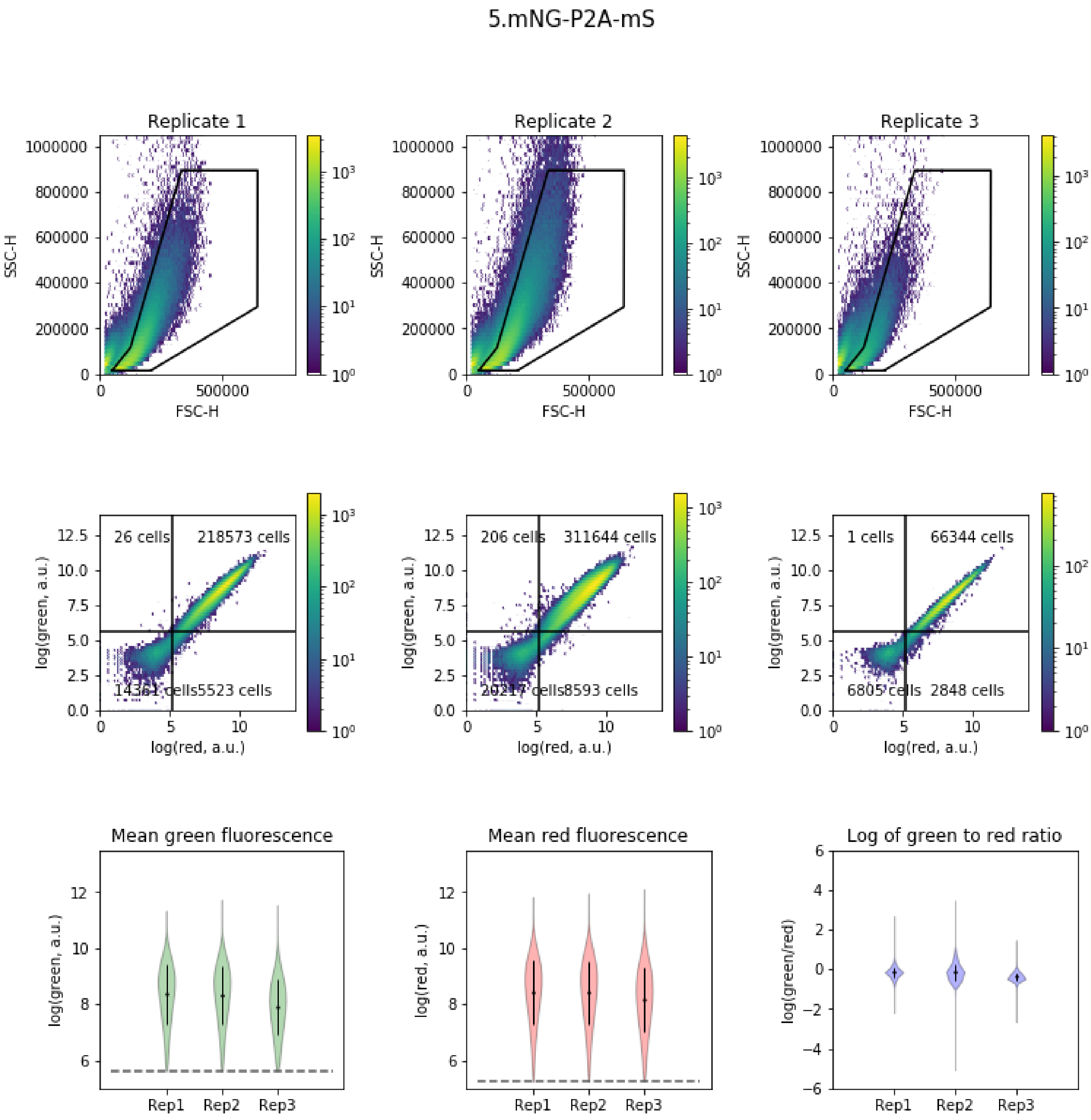


**Fig. S6. Flow cytometry data of AX4 cells transfected with mNeonGreen-P2A-mScarlet-I.Top.** Forward scatter and side scatter profiles of all three replicates. Density map color indicates event count. Events inside the polygon, defined as shown in S1, were considered live cells. **Middle.** Green and red fluorescence values for live cells for each replicate. Displayed numbers indicate total event count in a given quadrant. Cells in the upper right quadrant were retained for further analysis. **Bottom.** Mean fluorescence values and ratio of green to red fluorescence for each replicate. For each violin plot of single-cell values, the overlaid dot represents mean of the distribution and the error bars represent the standard deviation. Dashed lines represent fluorescence thresholds.

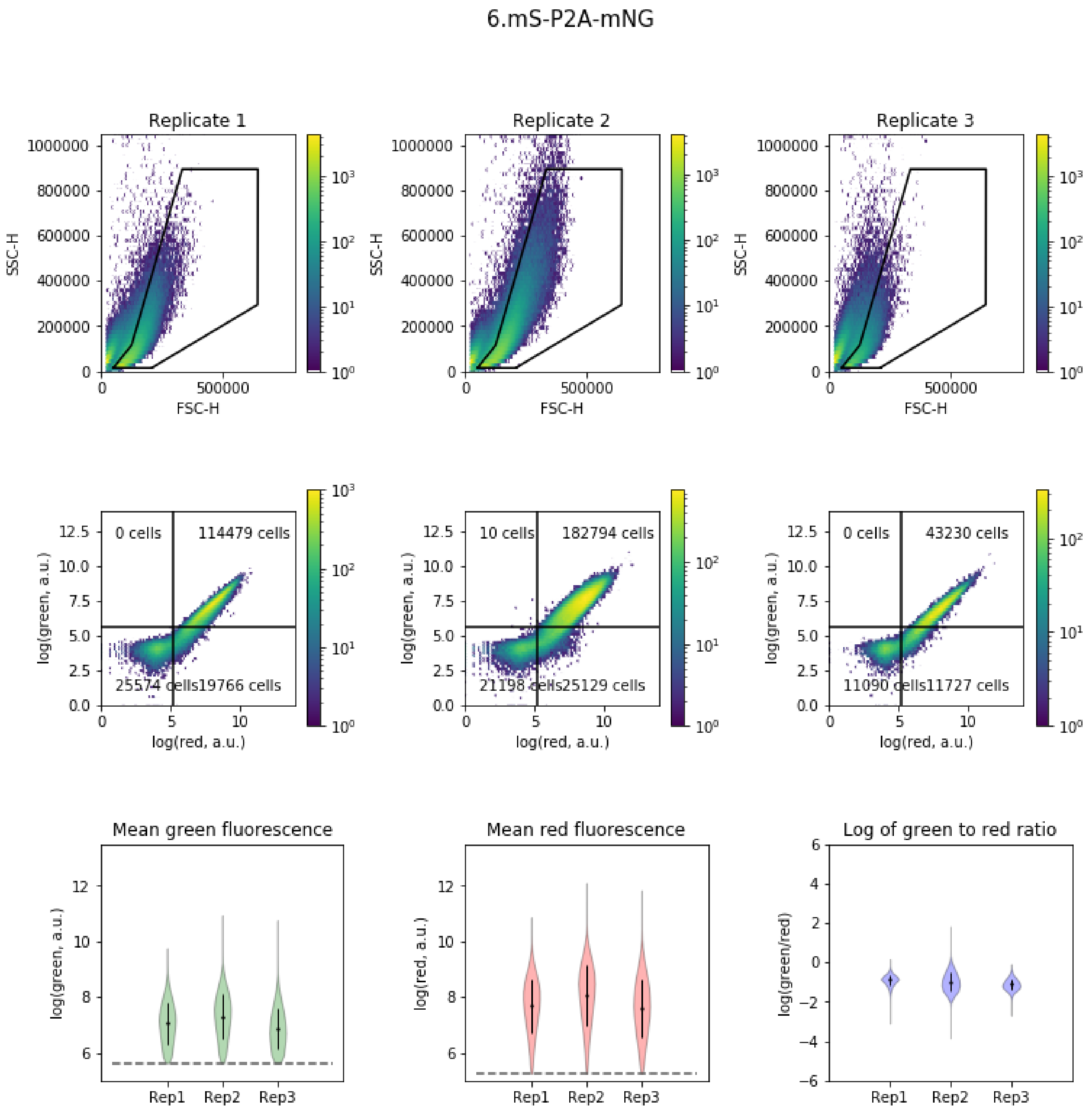


**Fig. S7. Flow cytometry data of AX4 cells transfected with mScarlet-I-P2A-mNeonGreen.Top.** Forward scatter and side scatter profiles of all three replicates. Density map color indicates event count. Events inside the polygon, defined as shown in S1, were considered live cells. **Middle.** Green and red fluorescence values for live cells for each replicate. Displayed numbers indicate total event count in a given quadrant. Cells in the upper right quadrant were retained for further analysis. **Bottom.** Mean fluorescence values and ratio of green to red fluorescence for each replicate. For each violin plot of single-cell values, the overlaid dot represents mean of the distribution and the error bars represent the standard deviation. Dashed lines represent fluorescence thresholds.

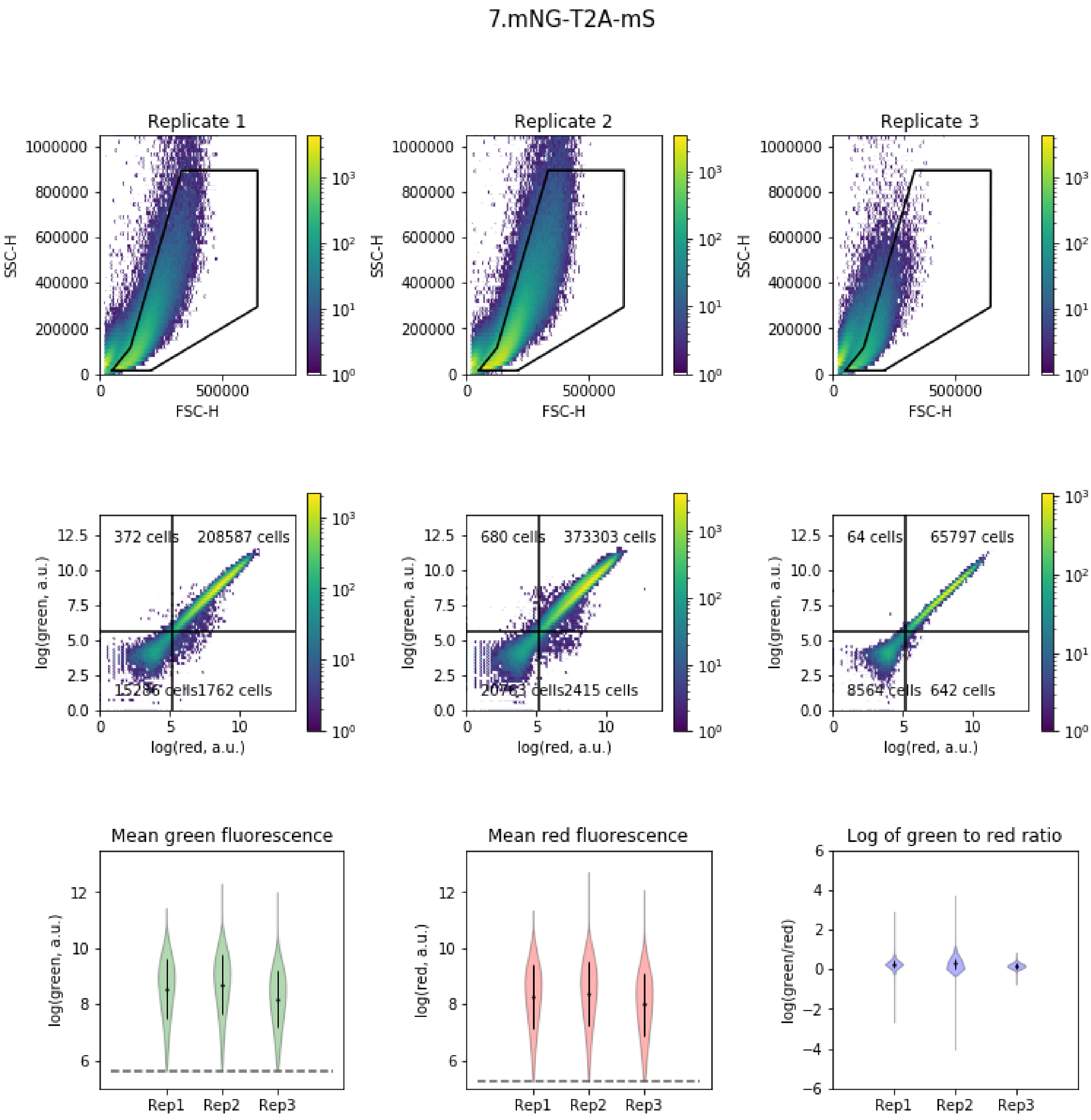


**Fig. S8. Flow cytometry data of AX4 cells transfected with mNeonGreen-T2A-mScarlet-I.Top.** Forward scatter and side scatter profiles of all three replicates. Density map color indicates event count. Events inside the polygon, defined as shown in S1, were considered live cells. **Middle.** Green and red fluorescence values for live cells for each replicate. Displayed numbers indicate total event count in a given quadrant. Cells in the upper right quadrant were retained for further analysis. **Bottom.** Mean fluorescence values and ratio of green to red fluorescence for each replicate. For each violin plot of single-cell values, the overlaid dot represents mean of the distribution and the error bars represent the standard deviation. Dashed lines represent fluorescence thresholds.

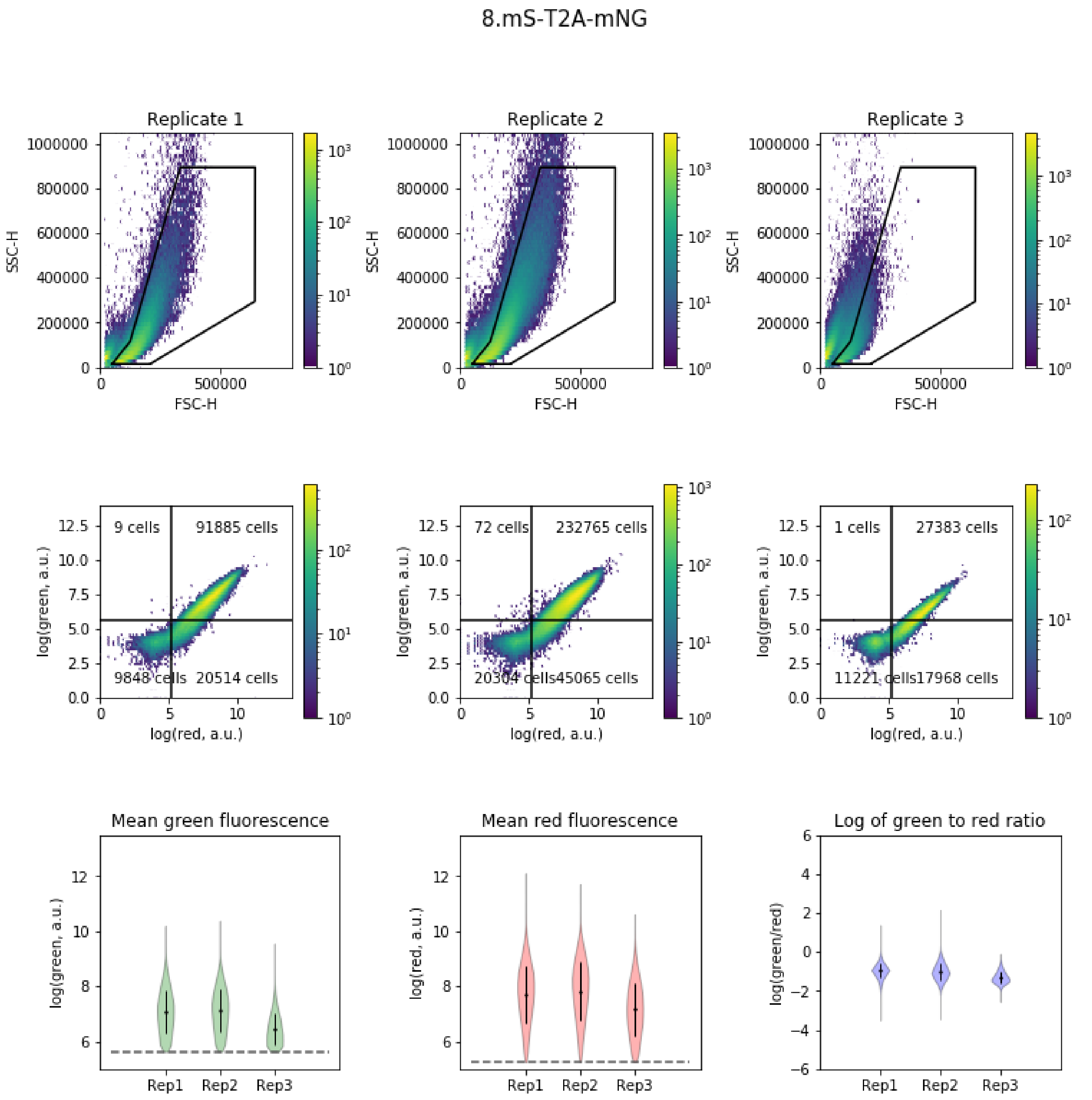


**Fig. S9. Flow cytometry data of AX4 cells transfected with mScarlet-I-T2A-mNeonGreen.Top.** Forward scatter and side scatter profiles of all three replicates. Density map color indicates event count. Events inside the polygon, defined as shown in S1, were considered live cells. **Middle.** Green and red fluorescence values for live cells for each replicate. Displayed numbers indicate total event count in a given quadrant. Cells in the upper right quadrant were retained for further analysis. **Bottom.** Mean fluorescence values and ratio of green to red fluorescence for each replicate. For each violin plot of single-cell values, the overlaid dot represents mean of the distribution and the error bars represent the standard deviation. Dashed lines represent fluorescence thresholds.

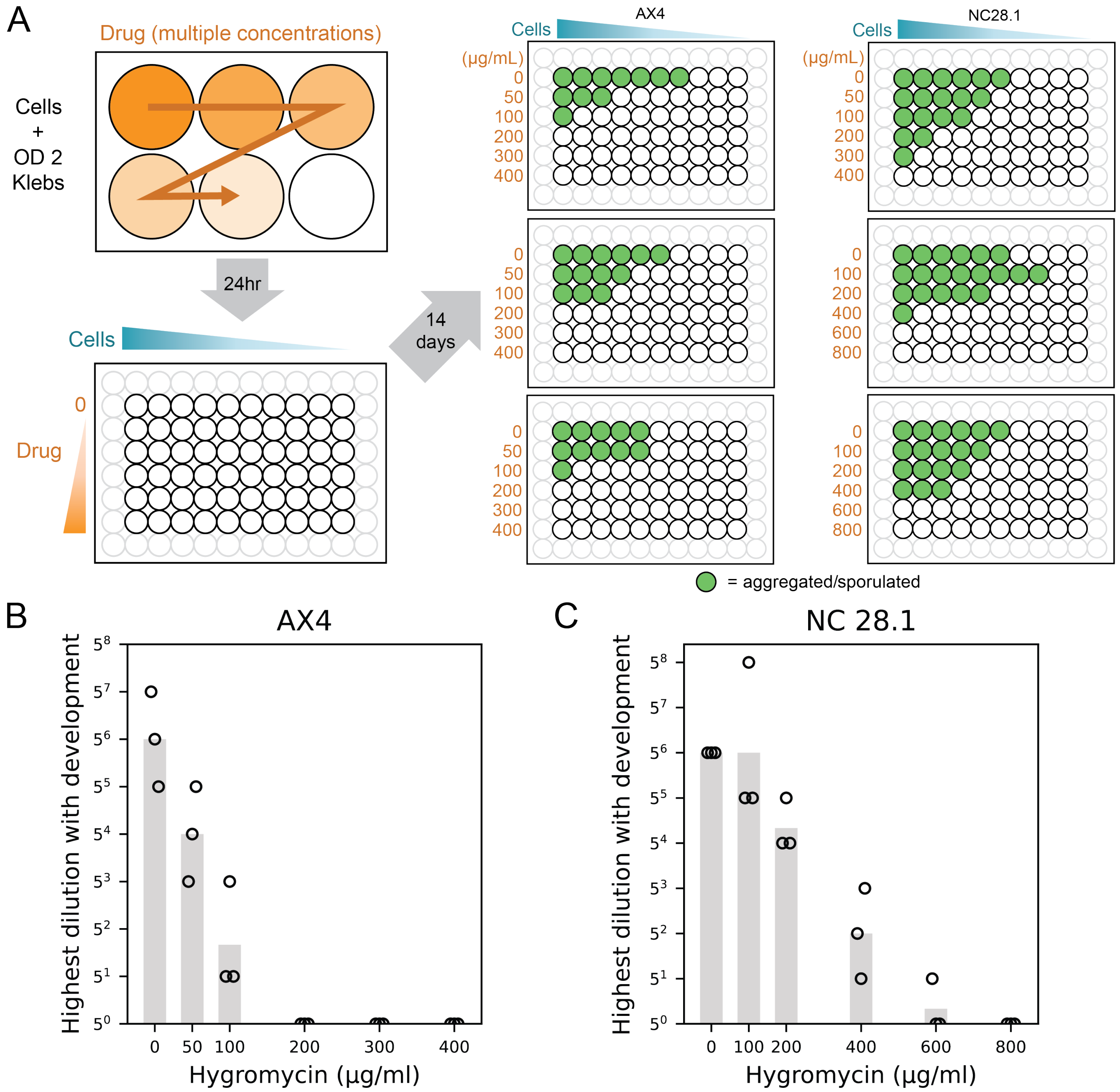


**Fig. S10. Assay of growth and development after treatment with hygromycin. A.** Diagram illustrating assay used to determine antibiotic susceptibility for different cell lines. Cells were grown with the indicated concentration of Hygromycin B Gold for 24 hours, then diluted and plated on SM agar. Wells were monitored for aggregation and development for two weeks after plating. At right, the final growth outcomes are shown for each replicate. **B-C.** Summarized assay results for AX4 (B) and NC28.1 (C). Each marker shows an independent experiment.

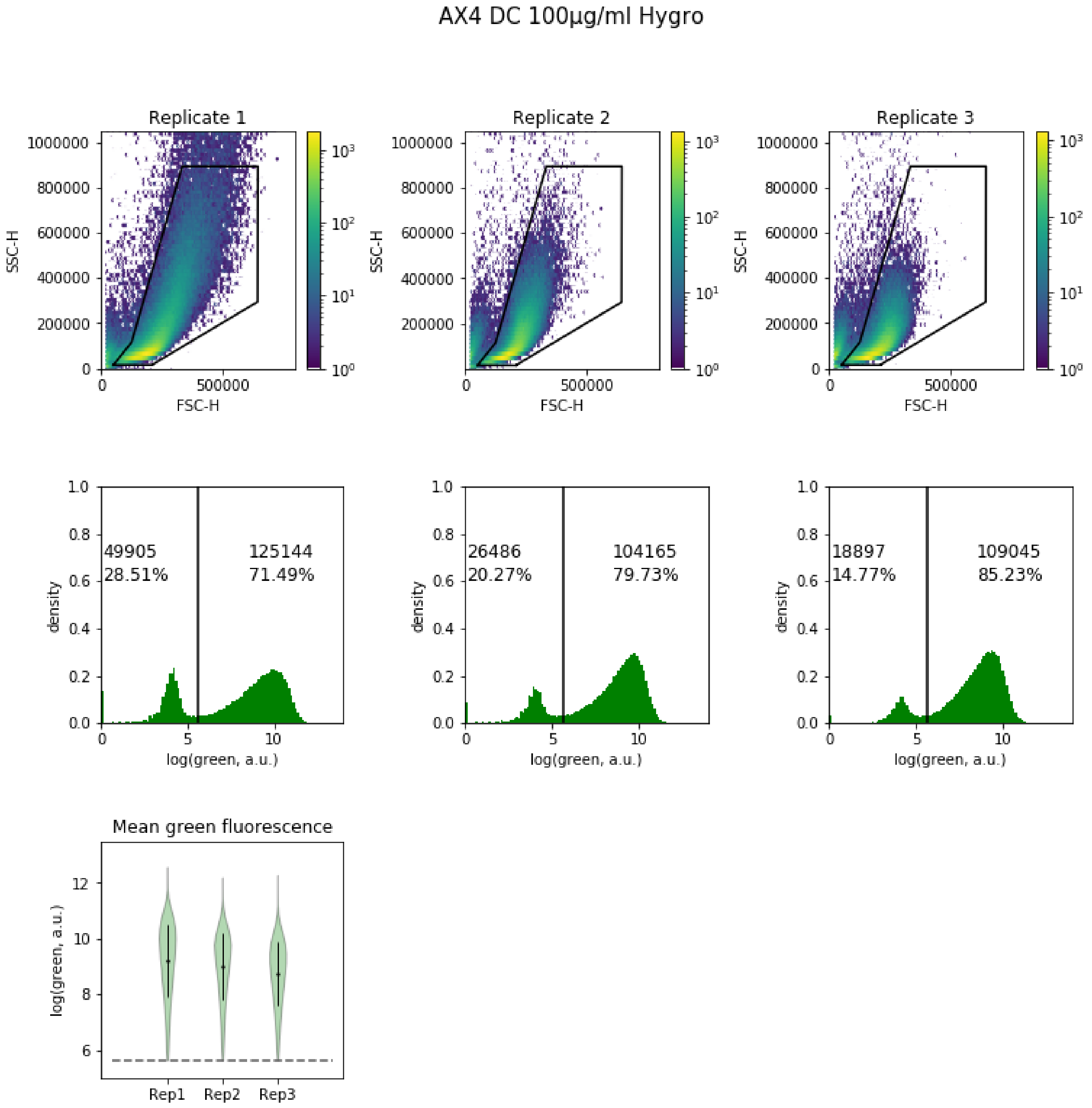


**Fig. S11. Flow cytometry data of AX4 cells transfected with the dual cassette plasmid and selected with 100 µg/ml hygromycin.Top.** Forward scatter and side scatter profiles of all three replicates. Events inside the polygon were considered live cells. **Middle.** Green fluorescence values for live cells for each replicate. Displayed numbers indicate the total event count and the percentage of events below or above the threshold. Cells above the fluorescence threshold were retained for further analysis. **Bottom.** Mean fluorescence values for each replicate.

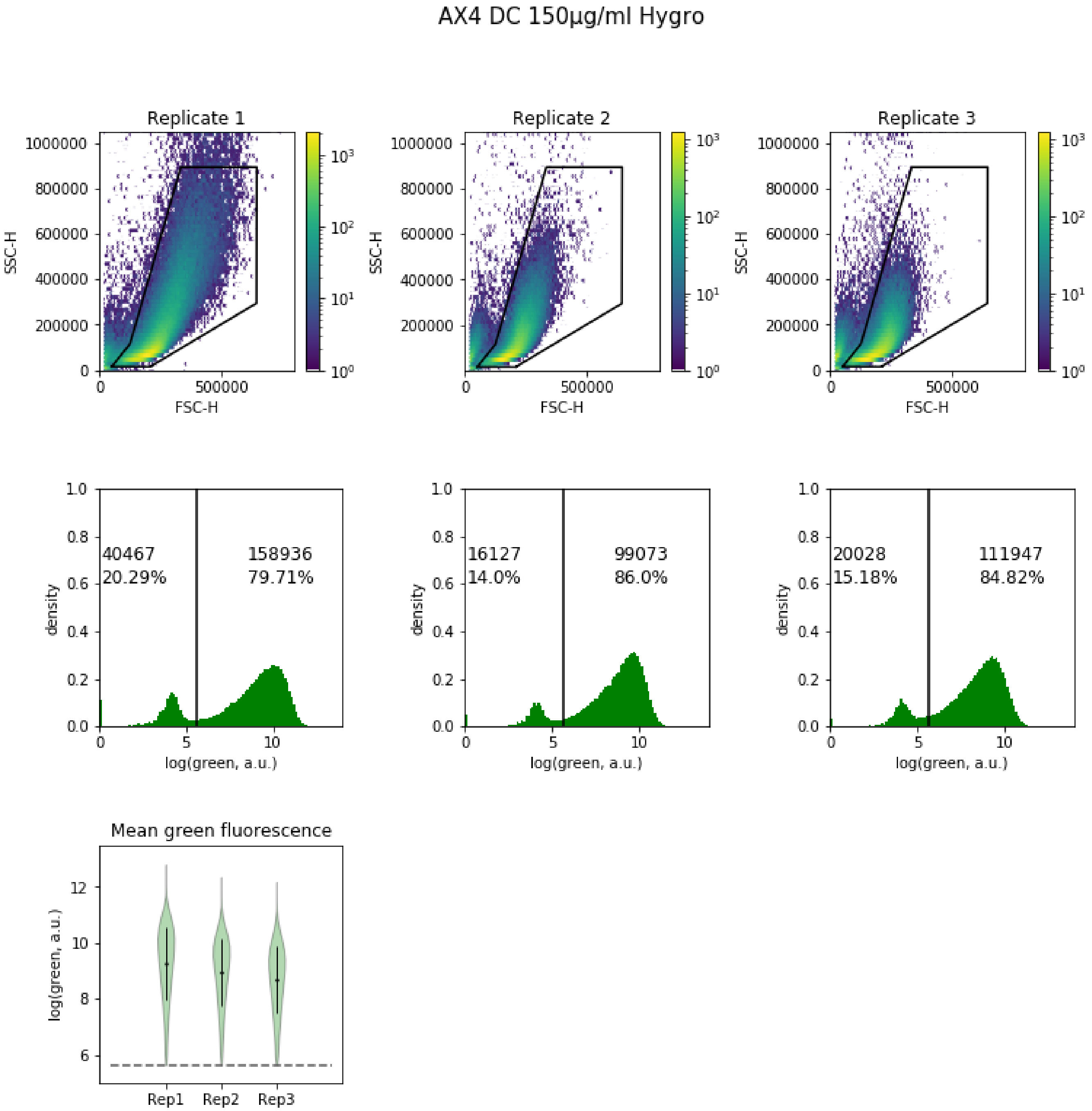


**Fig. S12. Flow cytometry data of AX4 cells transfected with the dual cassette plasmid and selected with 150 µg/ml hygromycin.Top.** Forward scatter and side scatter profiles of all three replicates. Events inside the polygon were considered live cells. **Middle.** Green fluorescence values for live cells for each replicate. Displayed numbers indicate the total event count and the percentage of events below or above the threshold. Cells above the fluorescence threshold were retained for further analysis. **Bottom.** Mean fluorescence values for each replicate.

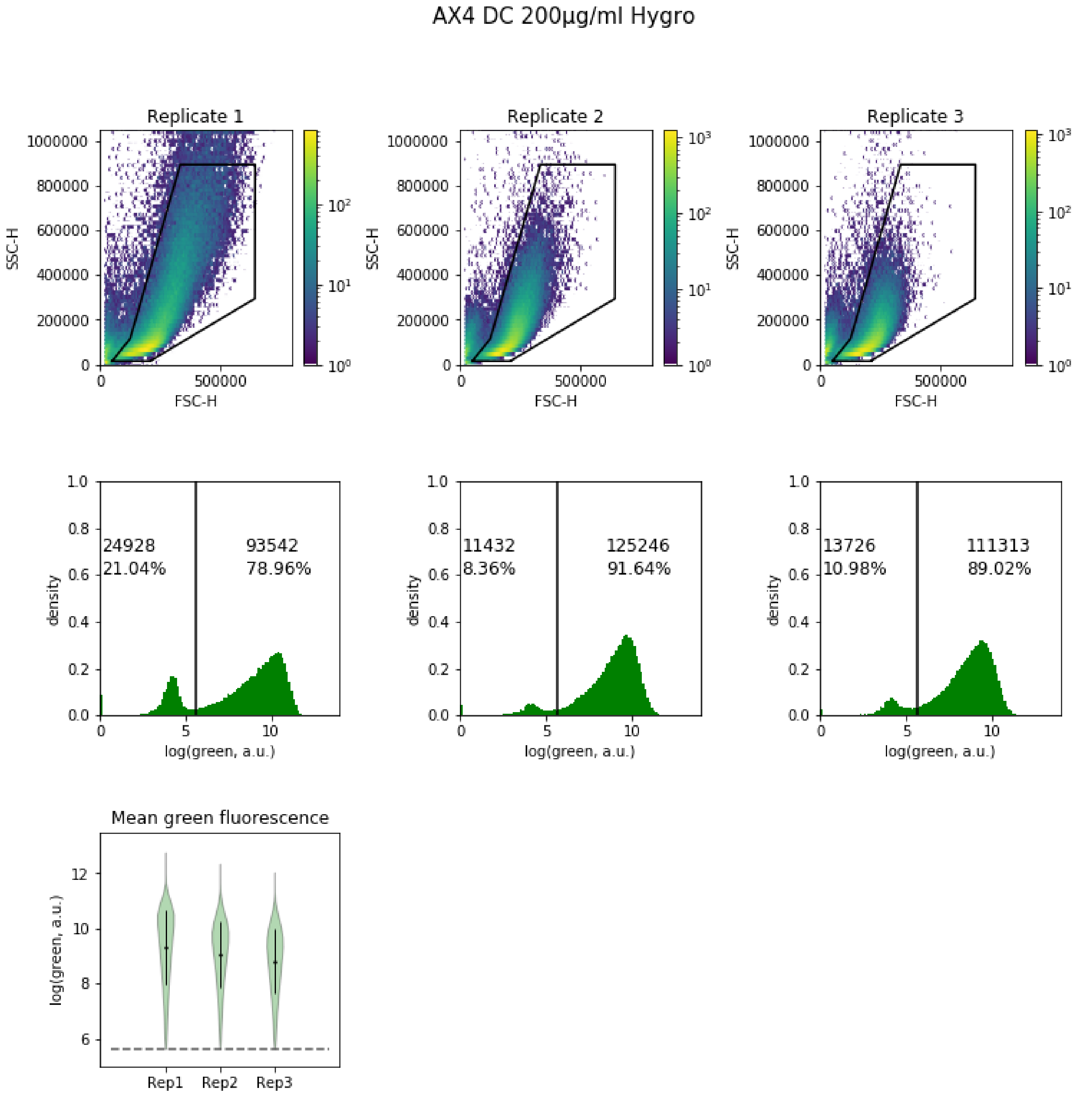


**Fig. S13. Flow cytometry data of AX4 cells transfected with the dual cassette plasmid and selected with 200 µg/ml hygromycin.Top.** Forward scatter and side scatter profiles of all three replicates. Events inside the polygon were considered live cells. **Middle.** Green fluorescence values for live cells for each replicate. Displayed numbers indicate the total event count and the percentage of events below or above the threshold. Cells above the fluorescence threshold were retained for further analysis. **Bottom.** Mean fluorescence values for each replicate.

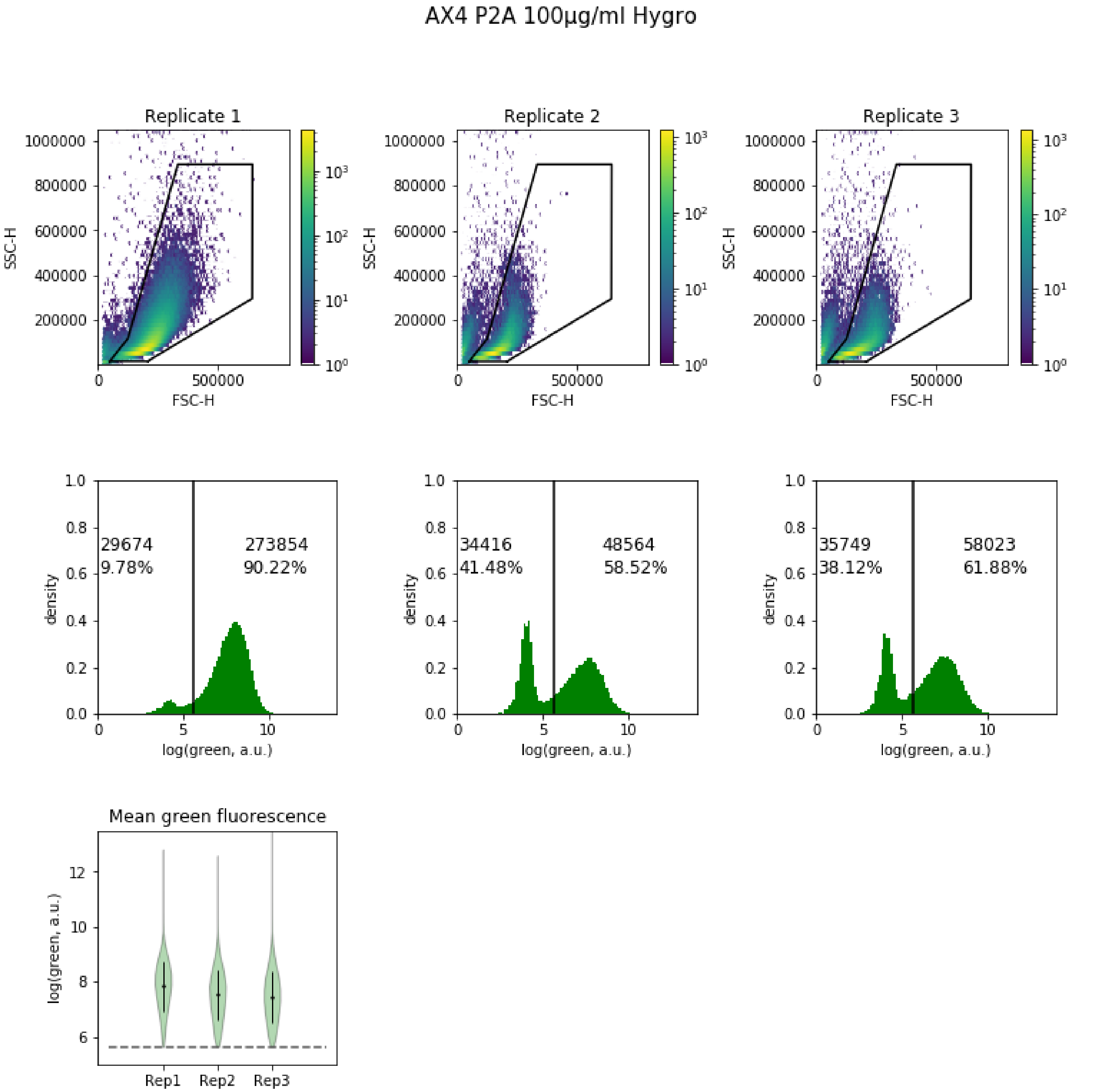


**Fig. S14. Flow cytometry data of AX4 cells transfected with the P2A plasmid and selected with 100 µg/ml hygromycin.Top.** Forward scatter and side scatter profiles of all three replicates. Events inside the polygon were considered live cells. **Middle.** Green fluorescence values for live cells for each replicate. Displayed numbers indicate the total event count and the percentage of events below or above the threshold. Cells above the fluorescence threshold were retained for further analysis. **Bottom.** Mean fluorescence values for each replicate.

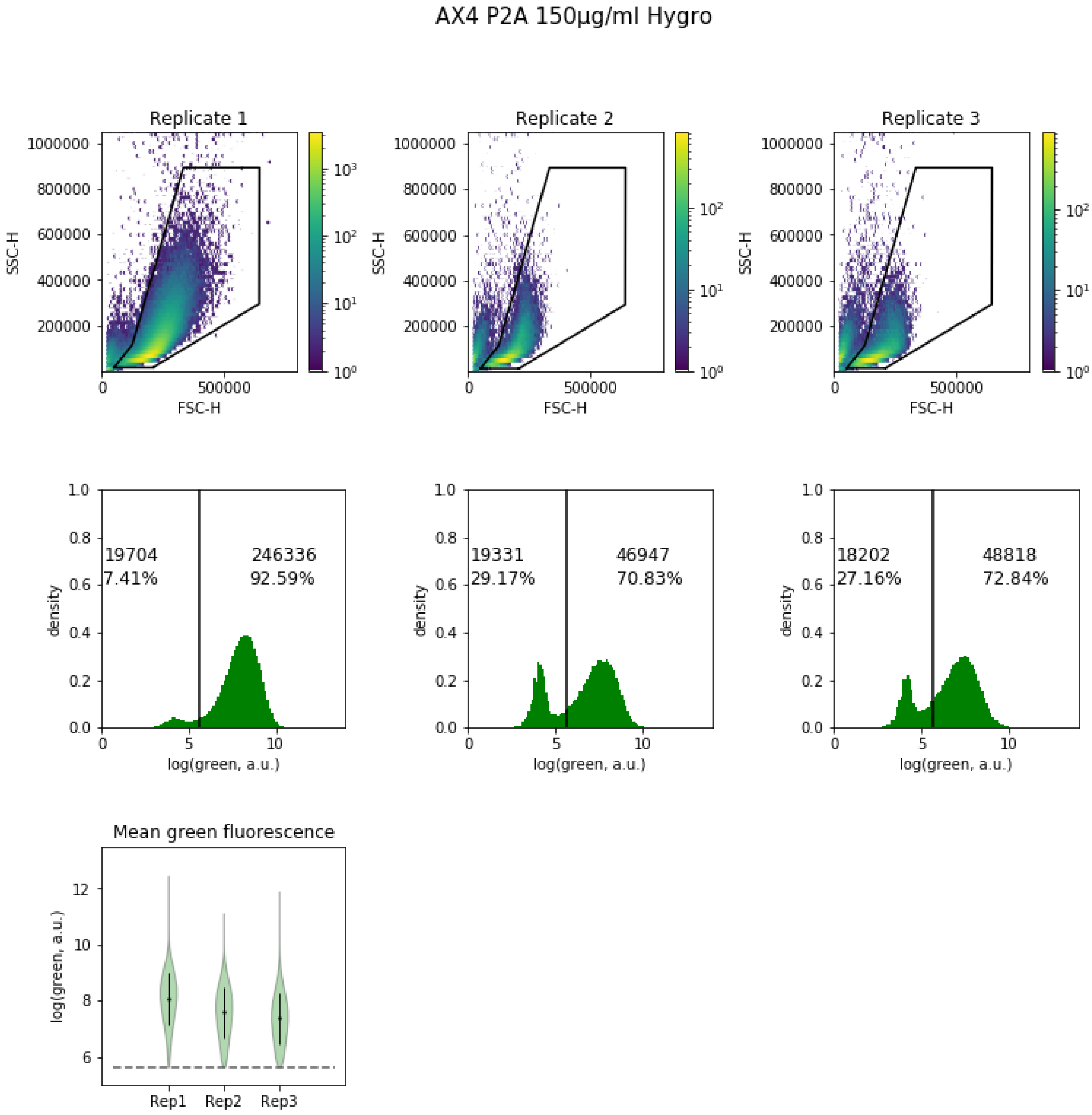


**Fig. S15. Flow cytometry data of AX4 cells transfected with the P2A plasmid and selected with 150 µg/ml hygromycin.Top.** Forward scatter and side scatter profiles of all three replicates. Events inside the polygon were considered live cells. **Middle.** Green fluorescence values for live cells for each replicate. Displayed numbers indicate the total event count and the percentage of events below or above the threshold. Cells above the fluorescence threshold were retained for further analysis. **Bottom.** Mean fluorescence values for each replicate.

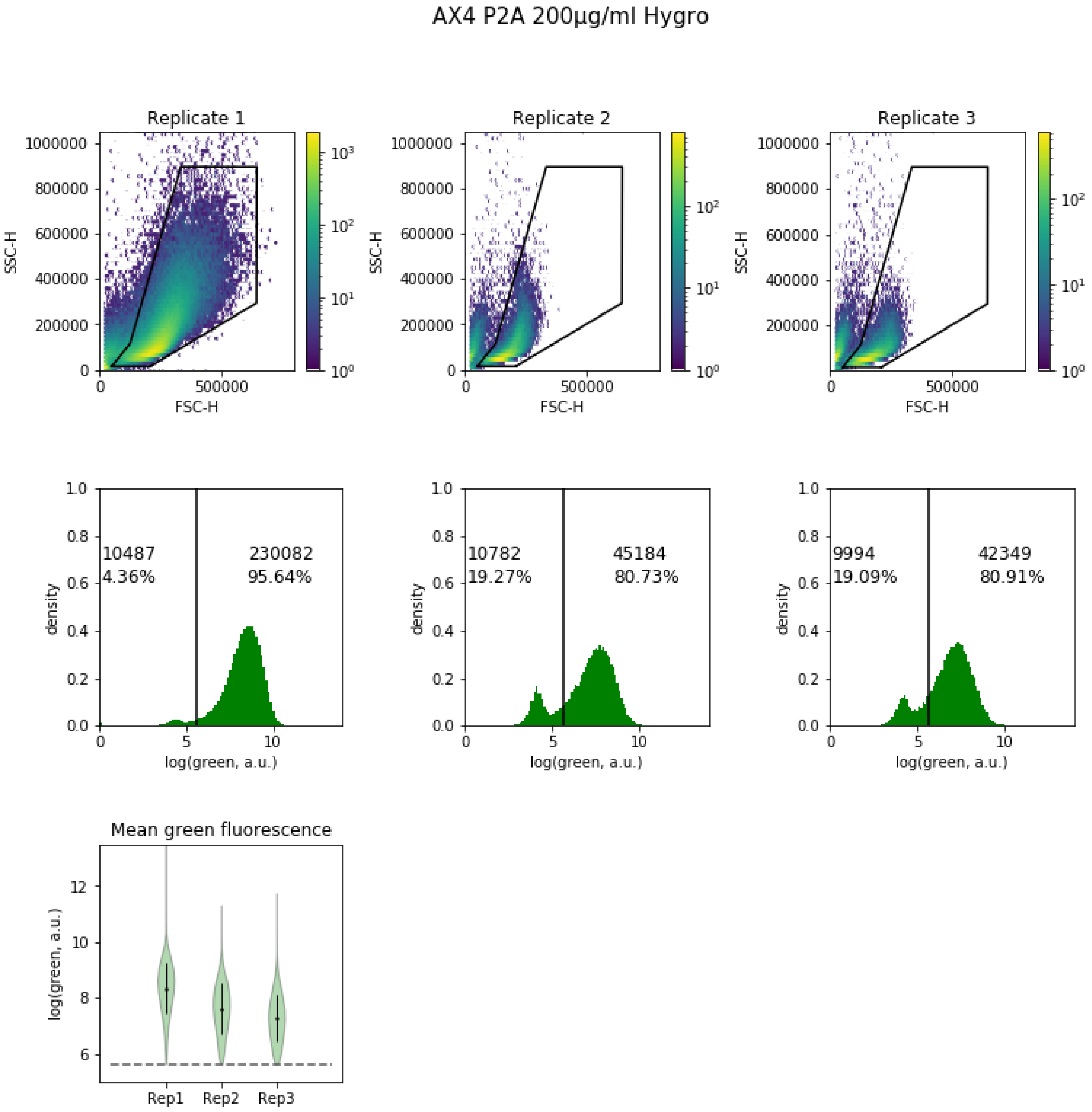


**Fig. S16. Flow cytometry data of AX4 cells transfected with the P2A plasmid and selected with 200 µg/ml hygromycin.Top.** Forward scatter and side scatter profiles of all three replicates. Events inside the polygon were considered live cells. **Middle.** Green fluorescence values for live cells for each replicate. Displayed numbers indicate the total event count and the percentage of events below or above the threshold. Cells above the fluorescence threshold were retained for further analysis. **Bottom.** Mean fluorescence values for each replicate.

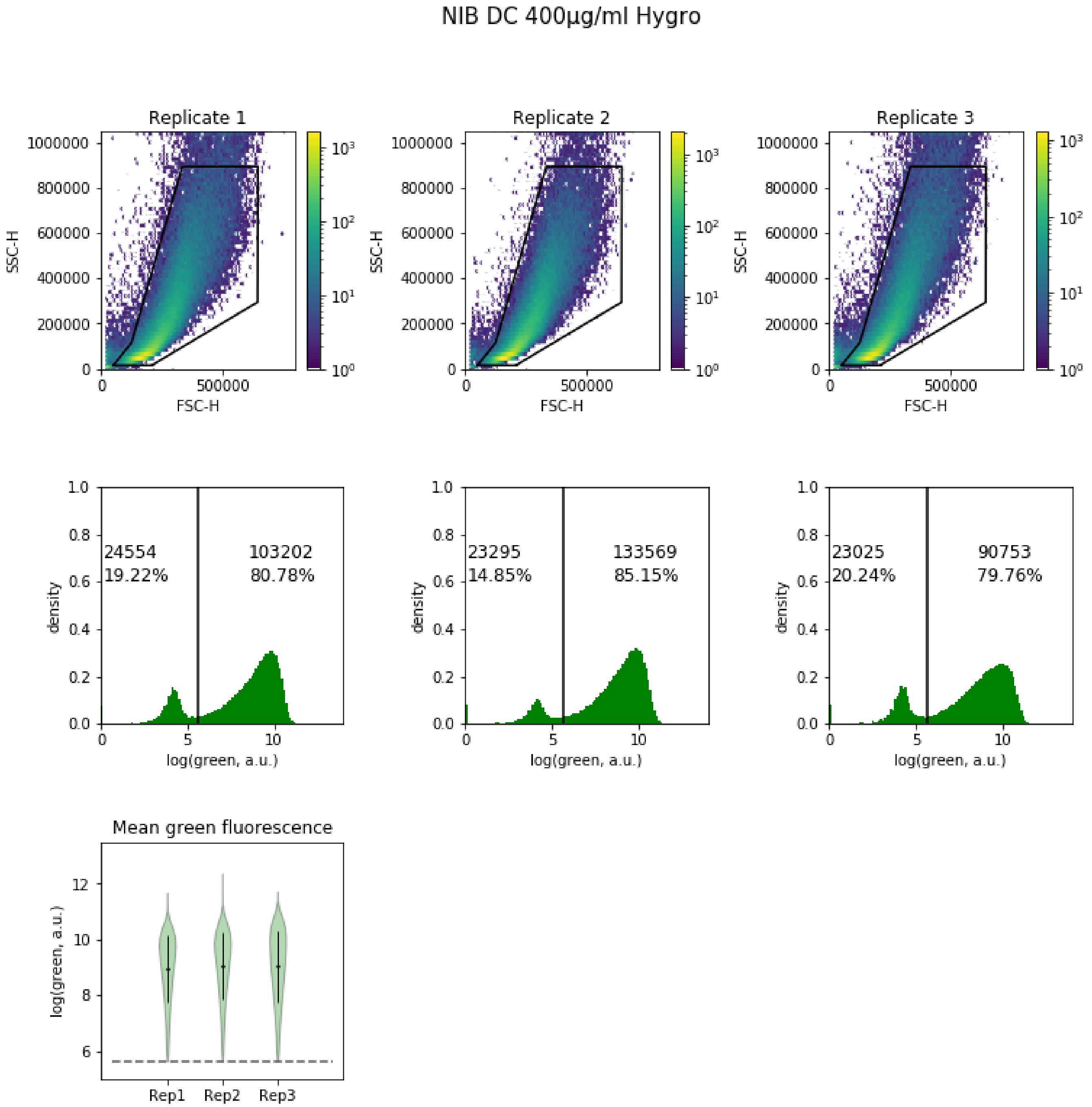


**Fig. S17. Flow cytometry data of NC28.1 cells transfected with the dual cassette plasmid and selected with 400 µg/ml hygromycin.Top.** Forward scatter and side scatter profiles of all three replicates. Events inside the polygon were considered live cells. **Middle.** Green fluorescence values for live cells for each replicate. Displayed numbers indicate the total event count and the percentage of events below or above the threshold. Cells above the fluorescence threshold were retained for further analysis. **Bottom.** Mean fluorescence values for each replicate.

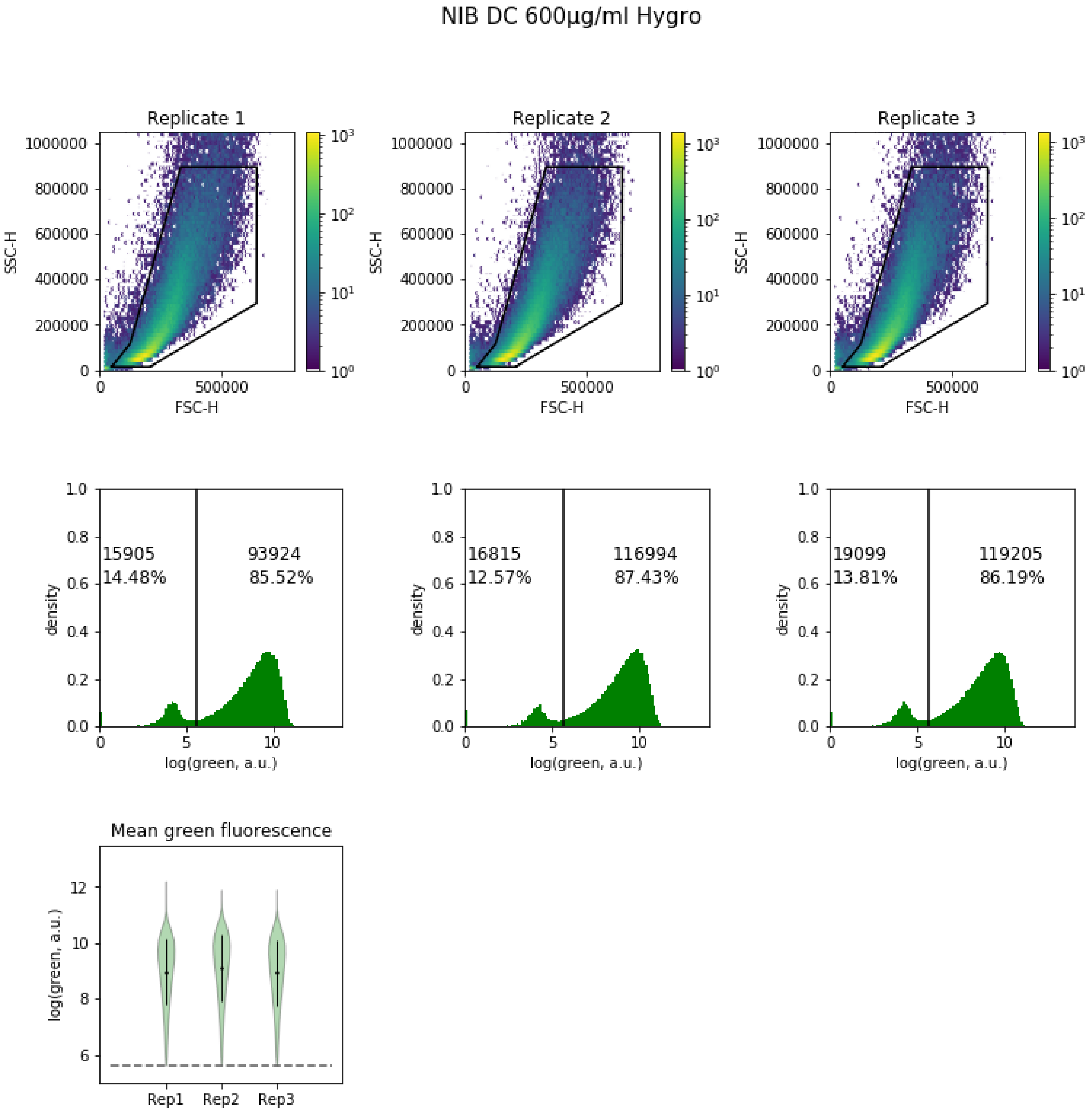


**Fig. S18. Flow cytometry data of NC28.1 cells transfected with the dual cassette plasmid and selected with 600 µg/ml hygromycin.Top.** Forward scatter and side scatter profiles of all three replicates. Events inside the polygon were considered live cells. **Middle.** Green fluorescence values for live cells for each replicate. Displayed numbers indicate the total event count and the percentage of events below or above the threshold. Cells above the fluorescence threshold were retained for further analysis. **Bottom.** Mean fluorescence values for each replicate.

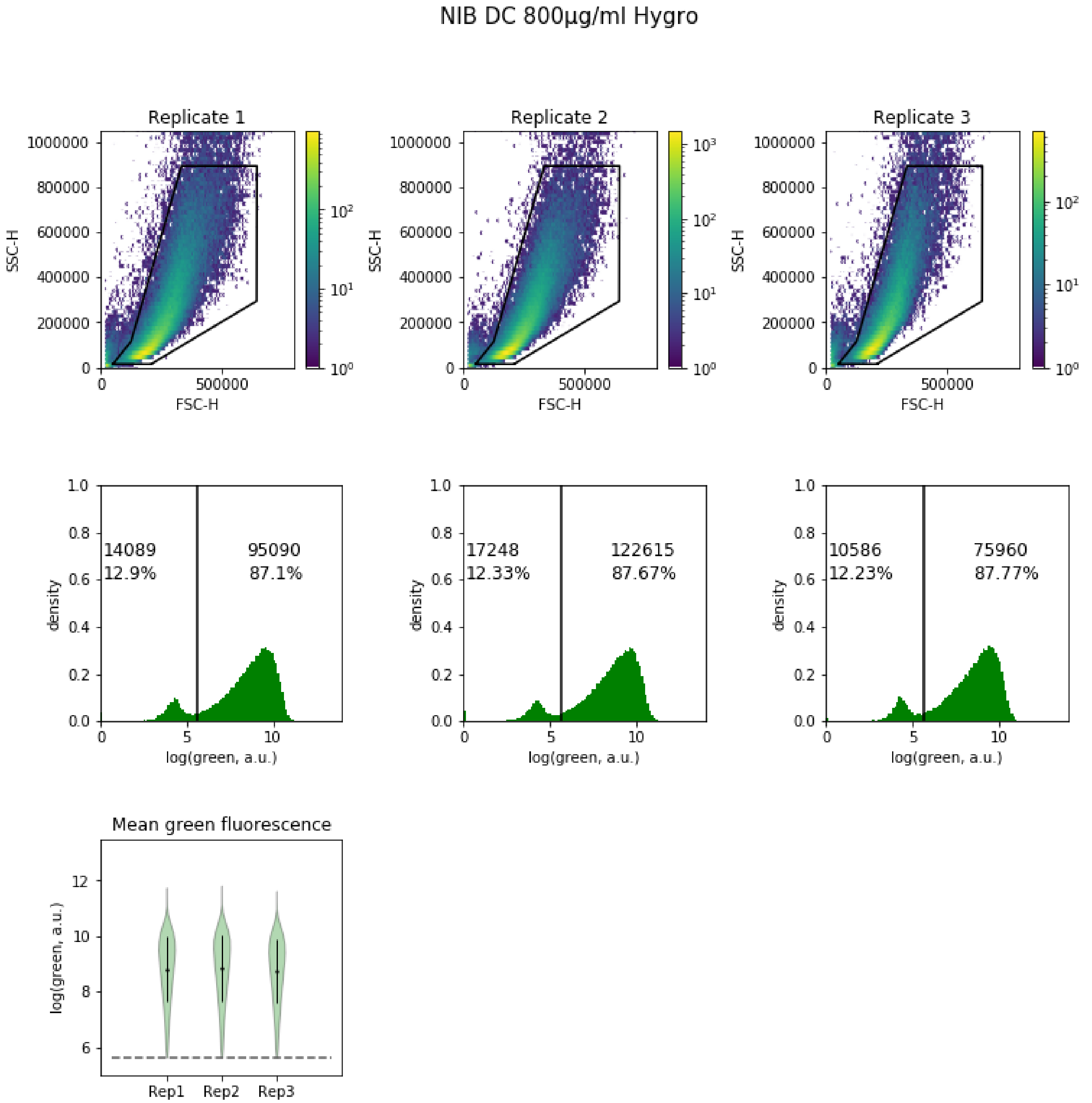


**Fig. S19. Flow cytometry data of NC28.1 cells transfected with the dual cassette plasmid and selected with 800 µg/ml hygromycin.Top.** Forward scatter and side scatter profiles of all three replicates. Events inside the polygon were considered live cells. **Middle.** Green fluorescence values for live cells for each replicate. Displayed numbers indicate the total event count and the percentage of events below or above the threshold. Cells above the fluorescence threshold were retained for further analysis. **Bottom.** Mean fluorescence values for each replicate.

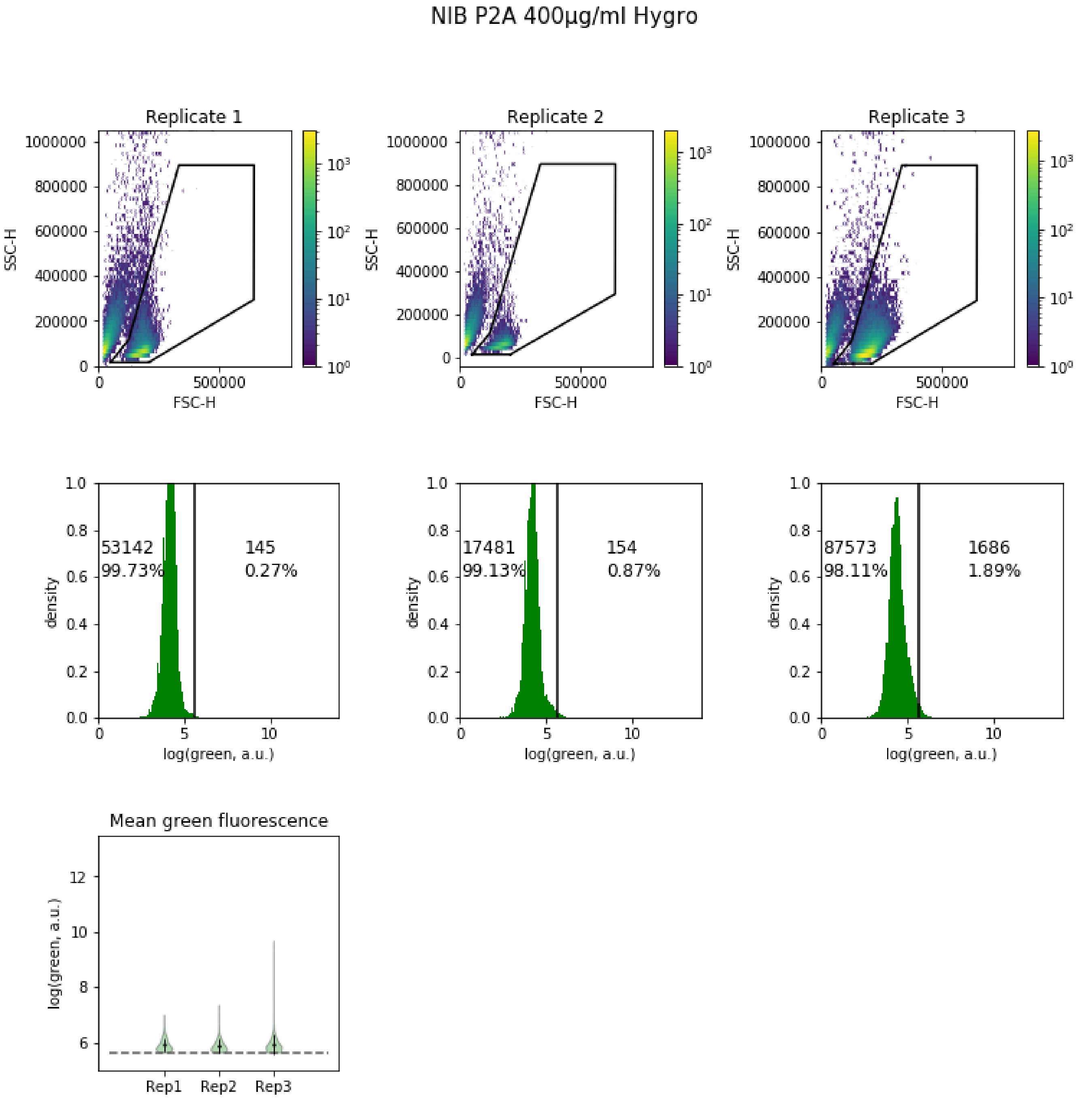


**Fig. S20. Flow cytometry data of NC28.1 cells transfected with the P2A plasmid and selected with 400 µg/ml hygromycin.Top.** Forward scatter and side scatter profiles of all three replicates. Events inside the polygon were considered live cells. **Middle.** Green fluorescence values for live cells for each replicate. Displayed numbers indicate the total event count and the percentage of events below or above the threshold. Cells above the fluorescence threshold were retained for further analysis. **Bottom.** Mean fluorescence values for each replicate.

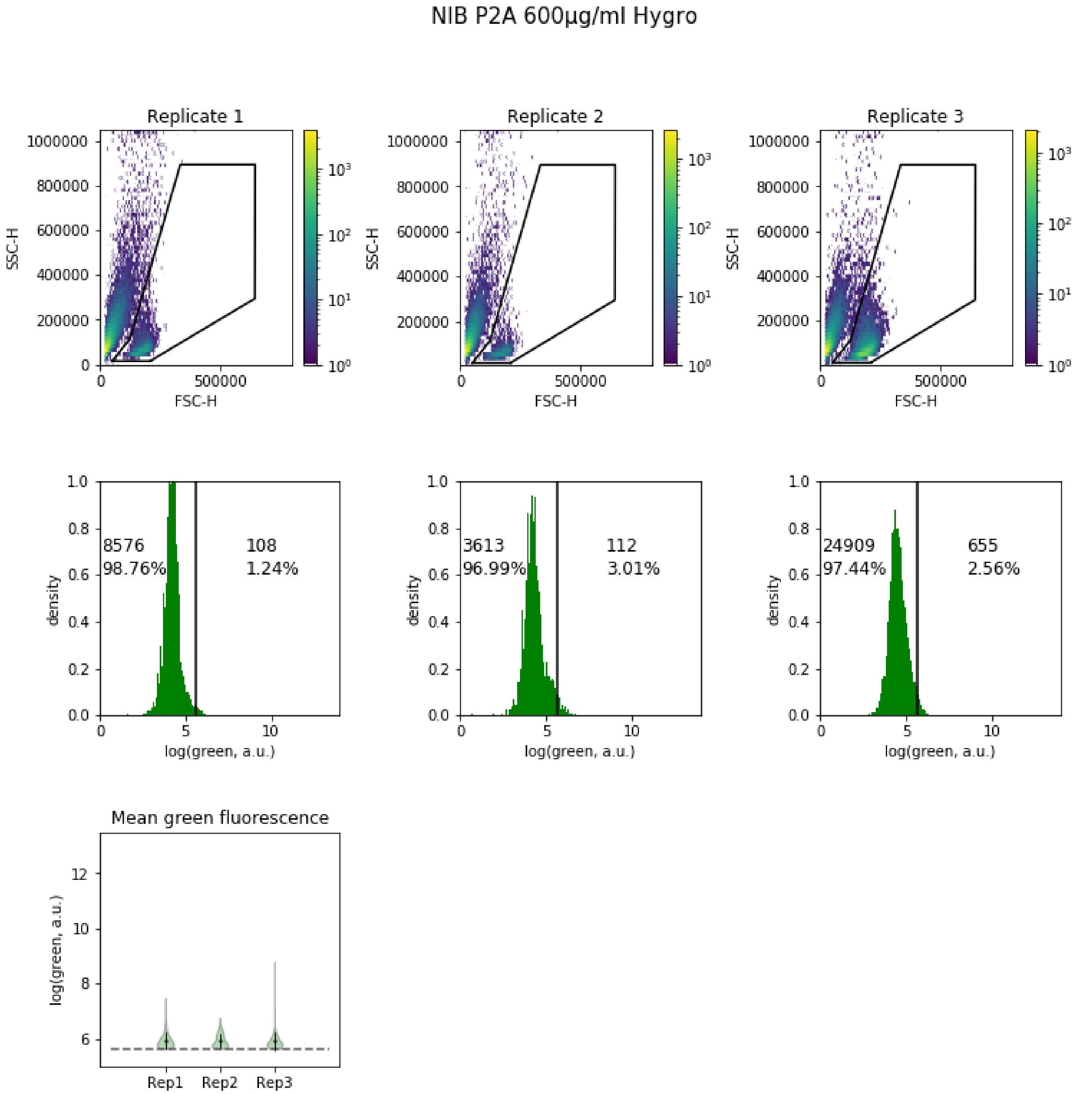


**Fig. S21. Flow cytometry data of NC28.1 cells transfected with the P2A plasmid and selected with 600 µg/ml hygromycin.Top.** Forward scatter and side scatter profiles of all three replicates. Events inside the polygon were considered live cells. **Middle.** Green fluorescence values for live cells for each replicate. Displayed numbers indicate the total event count and the percentage of events below or above the threshold. Cells above the fluorescence threshold were retained for further analysis. **Bottom.** Mean fluorescence values for each replicate.

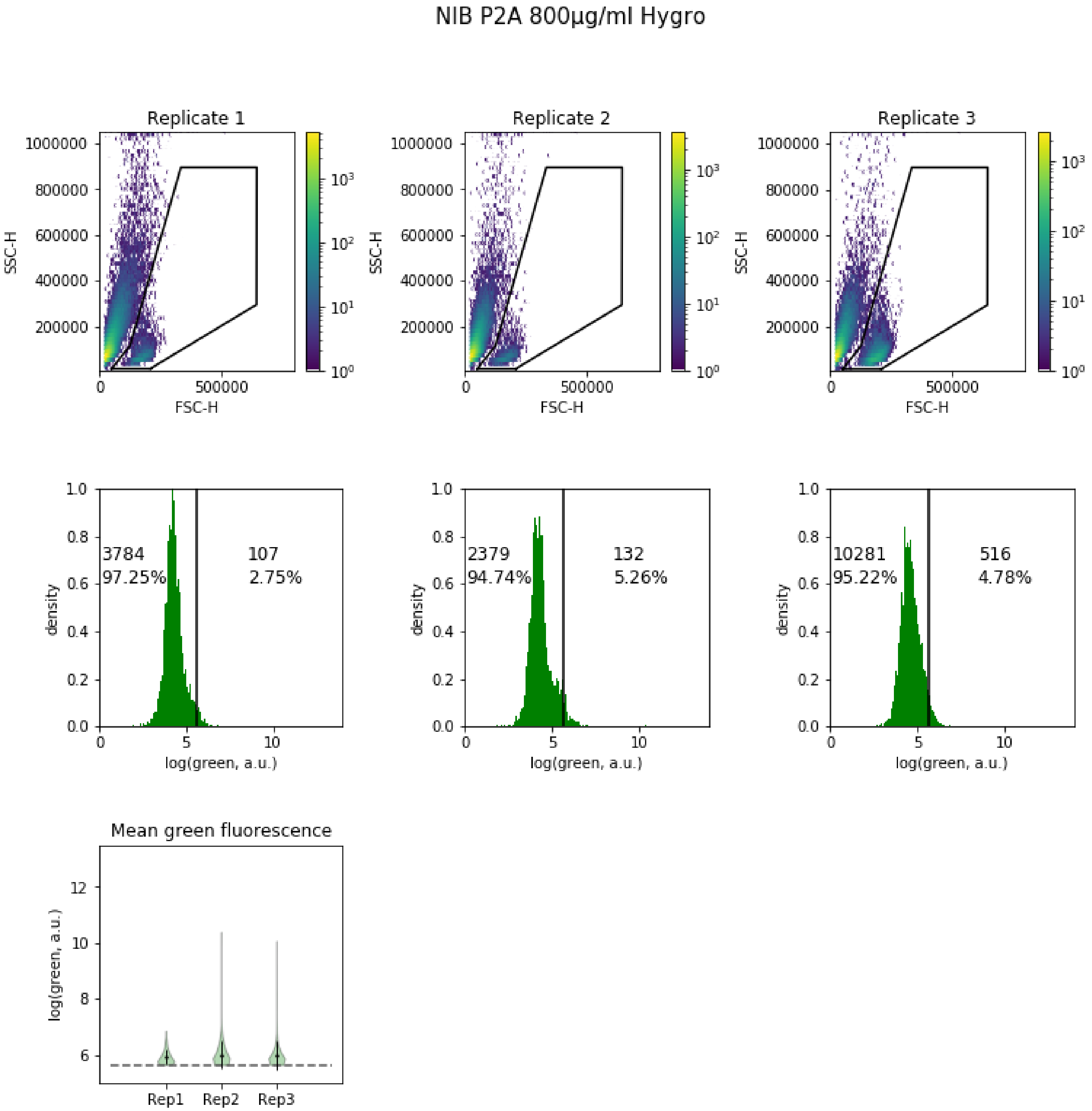


**Fig. S22. Flow cytometry data of NC28.1 cells transfected with the P2A plasmid and selected with 800 µg/ml hygromycin.Top.** Forward scatter and side scatter profiles of all three replicates. Events inside the polygon were considered live cells. **Middle.** Green fluorescence values for live cells for each replicate. Displayed numbers indicate the total event count and the percentage of events below or above the threshold. Cells above the fluorescence threshold were retained for further analysis. **Bottom.** Mean fluorescence values for each replicate.
